# Unbiased preclinical phenotyping reveals neuroprotective properties of pioglitazone

**DOI:** 10.1101/2024.08.30.610328

**Authors:** Edward C. Harding, Hsiao-Jou Cortina Chen, Dmytro Shepilov, Stefanie O. Zhang, Christine Rowley, Iman Mali, Jiahui Chen, Natasha Stewart, Dean Swinden, Sam J. Washer, Andrew R. Bassett, Florian T. Merkle

## Abstract

Animal models are essential for assessing the preclinical efficacy of candidate drugs, but animal data often fails to replicate in human clinical trials. This translational gulf is due in part to the use of models that do not accurately replicate human disease processes and phenotyping strategies that do not capture sensitive, disease-relevant measures. To address these challenges with the aim of validating candidate neuroprotective drugs, we combined a mouse prion (RML scrapie) model that recapitulates the key common features of human neurodegenerative disease including bona fide neuronal loss, with unbiased and machine learning-assisted behavioural phenotyping. We found that this approach measured subtle, stereotyped, and progressive changes in motor behaviour over the disease time course that correlated with the earliest detectable histopathological changes in the mouse brain. To validate the utility of this model system, we tested whether the anti-diabetic drug pioglitazone could slow prion disease progression. Pioglitazone crosses the blood-brain-barrier and has been shown to reduce neurodegenerative disease severity in other mouse models. We found that in addition to significantly slowing the emergence of early-stage clinical signs of neurodegeneration, pioglitazone significantly improved motor coordination throughout the disease time course and reduced neuronal endoplasmic reticulum stress. Together, these findings suggest that pioglitazone could have neuroprotective properties in humans, confirm the utility of the scrapie mouse model of neurodegeneration, and provide generalisable experimental and analysis methods for the generation of data-rich behavioural data to accelerate and improve preclinical validation.

## INTRODUCTION

Dementia remains one of the most significant unmet clinical needs facing an ageing world, with more than 70 million people currently affected and prevalence accelerating (Nichols *et al*., 2022; Livingston *et al*., 2024). Long term care is a significant economic burden, and existing therapies offer at best mild improvements in a subset of patients (Sims *et al*., 2023; Van Dyck *et al*., 2023). Radical, cost-effective preventative or mitigating interventions are required to reverse this trajectory.

Small molecule drugs have diverse secondary pharmacology that may be beneficial in conditions they were not originally indicated. Drug repurposing for brain disorders using compounds that are safe and well understood in humans remains the fastest approach for the discovery of new treatments. However, there are a number of challenges to efficiently identify and validate candidate compounds in preclinical *in vivo* models, where large-scale screening is impractical and prohibitively expensive. Approximately 9,000 small molecules are thought to be broadly safe for human administration, of which 4-5000 small molecules are approved for general clinical use (Gaulton *et al*., 2017; Mendez *et al*., 2019). Around 1000 of these have the chemical properties associated with penetration of the blood brain barrier (BBB) making them potentially suitable for targeting central nervous system (CNS) disorders (D. Segall, 2012; Wager *et al*., 2016). When considering which drugs to repurpose for neurodegenerative disease, the number of candidate drugs can be further reduced by eliminating classes of drugs, such as antipsychotics, that tend to be unsuitable due to their kinetics or adverse reactions, for an elderly target patient population (Mok *et al*., 2024). Whilst the prioritisation of drugs for repurposing for neurodegenerative conditions is also informed by *in vitro* screening and mechanistic studies, it still requires strong preclinical data from animal models. The quality of both the animal model, and the data from animal studies, are essential for increasing the likelihood of successful clinical trials.

Animal models commonly used in dementia research typically do not replicate all relevant features of human neurodegenerative disease. For example, some transgenic models based on familial mutations (e.g. 5xFAD) represent only a small subset of human cases, while over-expression models (e.g. APP^-NF(F)^) replicate only a subset of histopathological features (Sasaguri *et al*., 2017). Similarly, models combining rare variants and gene overexpression also have distinct aetiology and disease mechanisms than are seen in the majority of human patients. Furthermore, use of these transgenic lines requires expensive and time-consuming breeding paradigms that complicate pre-clinical validation (Voelkl *et al*., 2020). In contrast, mouse models of prion disease accurately phenocopy human prion disease (Watts and Prusiner, 2014), and undergo a stereotyped disease progression combining clear behavioural deficits such as progressive loss of motor coordination with the full suite of histopathological hallmarks of human neurodegeneration, including synaptic dysfunction and loss, endoplasmic reticulum (ER) stress, neuroinflammation, and neuronal loss (Mallucci, 2009; Watts and Prusiner, 2014). Prion models can be generated on any genetic background, and the use of strains such as Rocky Mountain Laboratory (RML) scrapie, that normally infects sheep and does not cross the species barrier into humans (Galassi, Henneberg and Rühli, 2016), makes them a powerful and tractable model for preclinical studies.

Mouse models of neurodegenerative disease are typically phenotyped using a combination of behavioural scoring in standardised tests (e.g. rotarod, novel object recognition) and brain histology. Recently, advances in pose-estimation methods for animal tracking have made the extraction of feature-rich datasets from video recordings both practical and highly reliable (Mathis *et al*., 2018; Pereira *et al*., 2022) while the downstream application of machine learning methods to tracking data has allowed classification and phenotyping of animal behaviour (Biderman *et al*., 2023; Tillmann *et al*., 2024). This has been accompanied by advances in unsupervised time-series analysis (Hsu and Yttri, 2021) and the use of variational autoencoders and semi-supervised methods for detecting motor sequences or motifs from tracking data (Wiltschko *et al*., 2020; Luxem *et al*., 2022; Weinreb *et al*., 2024).

These methods have not yet been widely applied to systematic preclinical drug discovery but represent an exciting opportunity to test the potential neuroprotective role of drugs that target metabolic disease, which has been suggested to share mechanisms with neurodegenerative disease (Craft, 2009).

Extensive observational data has suggested that effective treatment of diabetes reduces dementia risk (Whitmer, 2009; Biessels *et al*., 2014; Samaras *et al*., 2020; Chen *et al*., 2023). Our own work on metformin demonstrated direct evidence of neuroprotection separate from its anti-diabetic mechanism (Harding *et al*., 2023), but it is unclear if these results extend to other anti-diabetic drugs with different molecular mechanisms. We considered multiple drug candidates for this work and chose to focus on molecules that are well tolerated in the elderly patient population, likely to cross the BBB, and are orally bioavailable. Pioglitazone is one such compound and also a second-line treatment for diabetes. Often used together with metformin, it has a different mechanism of action as an agonist on the PPARγ receptor. This allows us to test the hypothesis of generalisable neuroprotection from this class of compounds without the involvement of the AMPK target of metformin.

Here we show a preclinical pipeline using neural network-based pose estimation combined with traditional machine learning to speed up compound testing and validation. This approach provides unbiased quantification of motor signs and treatment efficacy, in mixed sex groups. Further, we show that these systems are more sensitive than traditional methods, detecting disease at an earlier time point and with less intra-animal variability.

Finally, we show that this model can provide an unbiased detection of the potential neuroprotective effects of pioglitazone at earlier time points than is achievable with traditional methods. Overall, this system is designed to generalise to any mouse model with motor deficits and can be used efficiently for preclinical phenotyping at scale.

## RESULTS

### Scrapie-inoculated mice have stereotyped signatures of disease progression

To establish a scrapie mouse model for neuroprotective drug testing, we first characterised the time course and signs of neurodegenerative disease progression in our laboratory, since these can be modulated by genetic background, sex, and prion strain and preparation methods (Morales, Abid and Soto, 2007; Akhtar *et al*., 2011). We therefore unilaterally inoculated the brains of 4-week old C57Bl6/J mice with either normal brain homogenate (NBH) or brain homogenate from mice carrying the RML strain of scrapie and characterised the time course of disease progression using traditional phenotyping approaches (Fig. 1A). We found that mice inoculated with RML lost a significant fraction of their body weight relative to controls starting around 14 weeks post-inoculation (wpi) (Fig. 1B), as previously described (Harding *et al*., 2023). We also found that RML mice started displaying overt signs of neurodegenerative disease starting at 17 wpi (Fig. 1C), including progressive loss of motor function that affected nearly all mice by week 20 (Fig. 1D). We corroborated these findings by recording the locomotor behaviour of NBH and RML mice and found that they displayed distinct locomotor behaviour by week 20, with RML mice showing hyperactivity and corner preferences (Fig. 1L and Supp. Fig. 1C), and turning behaviour (Supp. Fig. 1D).

**Figure 1.**
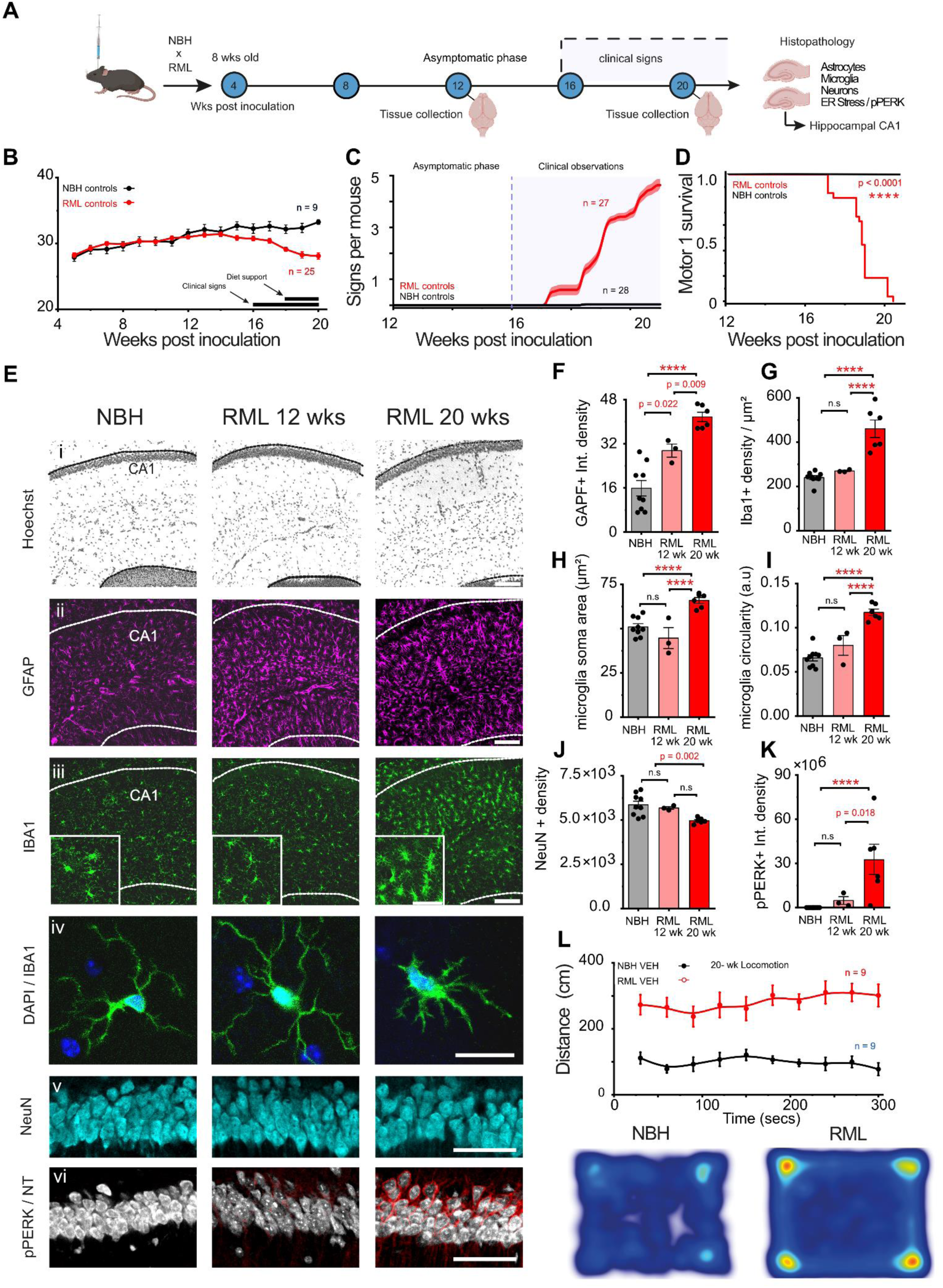
Time course of scrapie pathogenesis from asymptomatic to clinical signs by 20 weeks. (A) Experimental paradigm showing scrapie inoculation in 4-6 week old mice. Brains were collected at 12 and 20 weeks post inoculation (wpi) for histological analysis. (B) Mouse body weight in control (n=9) and scrapie mice (n=20) diverges from 12 weeks to 20 wpi and the trajectory is not altered by provision of wet food as diet support. (C) Clinical signs start around 17 wpi and rapidly accumulate providing a clear distinction between control (n = 28) and scrapie inoculated mice (n = 27). (D) Time to motor survival 1 (M1 *see methods*) starts at 17 wpi and is present in all scrapie inoculated mice by 21 wpi. (E) Histology panel for hippocampal CA1 region showing immunostaining for markers of microglia (Iba1), astrocytes (GFAP), neurons (NeuN) and ER stress (pPERK and NeuroTrace). Scale bars for respective images (i-iii) are 100 μm, (iv) is 20 μm, (v & vi) is 50 μm. (F) Integrated density of GFAP positive astrocytes per mm^2^. (G) Density of Iba1 positive microglia per mm^2^. (H) Iba1 positive soma area. (I) Iba1 cell circularity. (J) NeuN positive neuronal density per mm^2^ of CA1 pyramidal layer. (K) Integrated density of pPERK in pyramidal neurons per mm^2^. (L) 5-min locomotion assay showing differences in movement between control (n = 9) and scrapie inoculated mice (n = 9) and differences in thigmotaxis (heatmaps). Error bars are mean ± S.E.M. P values are shown on the graphs unless < 0.0001, which are represented as ****.

To correlate these physiological and behavioural changes with cellular changes in the brain, we performed immunohistochemistry on the brains of NBH- and RML-injected mice prior to the onset of overt behavioural phenotypes (12 wpi) and also after these phenotypes were present in the vast majority of RML mice (20 wpi). We focused on the CA1 region of the hippocampus contralateral to the injection site, since it is both close to the site of injection and clinically well associated with pathology in common dementias such as Alzheimer’s disease (Fig. 1A). Example histology is shown in Fig. 1E (i - vi). We observed an increase in immunoreactivity for glial fibrillary acidic protein (GFAP) by 12 wpi and 20 wpi (Fig. 1F), suggesting increased astrocyte reactivity over the course of disease progression (Smith *et al*., 2020). These histological changes were accompanied by the increased immunoreactivity for the microglial marker IBA1 (Fig. 1G), as well as increased microglial soma size (Fig. 1H) and circularity (Fig. 1I) in RML mice, suggesting microglial activation (Clarke, Crombag and Hall, 2021; Woodburn, Bollinger and Wohleb, 2021). High magnification images of microglia are shown in Fig. 1E (iv) with separate channels in Supp. Fig. 1A.

Since prion-induced proteostatic stress is associated with ER stress, we immunostained for the ER stress marker pPERK (Hughes and Mallucci, 2019). We found that while pPERK levels were virtually undetectable in control animals, they were significantly higher in RML mice at 12 wpi and highly elevated by 20 wpi, indicating early and ongoing ER stress (Fig. 1K). This neuronal ER stress was accompanied by the modest but significant loss of NeuN-positive cells from the CA1 region of the hippocampus at 20 wpi (Fig. 1J), indicating neuronal loss. Separate channels are shown in Supp. Fig. 1B.

Overall, these results are consistent with previous reports of progressively increasing inflammation associated with misfolding of the PrP^sc^ that results in neuronal ER stress, synaptic loss and eventually death (Hughes and Mallucci, 2019). Specifically, the time course of these changes in C57Bl/6J mice confirms the utility of the RML scrapie model for studying the full range of pathological features associated with many human neurodegenerative diseases (Freeman and Mallucci, 2016).

### Unbiased machine learning approaches can accurately classify scrapie disease progression

Classical assessment of ‘survival’ in neurodegenerative models including scrapie is based on the observed onset of early and/or confirmatory clinical signs, allowing disease endpoints to be determined before animals actually die. In addition to being more ethically sound, scoring signs related to motor impairment can help stage disease initiation and progression. The most common confirmatory sign in mice inoculated with RML scrapie is the impaired righting reflex, resulting largely from a one-sided weakness (contralateral to the inoculation site) in back-paw strength and coordination. This is preceded by the progressive loss of motor coordination that is sometimes detected as an early sign. However, it is challenging to consistently score complex movement, and standardisation relies heavily on the expertise and consistency of a trained observer. User-dependent and lab-dependent variation, along with high behavioural workloads and the challenges of group blinding (and unintended unblinding), create challenges for both the collection and interpretation of this type of data.

To implement an automated clinical assessment of mice as they progressed through the clinical stages of scrapie, we recorded approximately 2500 videos of mouse behaviour at 12, 16, and 20 wpi in a purpose-built frame designed to maintain consistent recording across different modalities (Figure 2A). To accurately capture variation in mouse movement, we recorded in three environments: an open field for top-down motion (Fig. 2B), on a clear platform for bottom-up gait analysis (Fig. 2C), and on a metal mesh to capture motor impairment and paw slipping (Fig. 2D). Recordings were designed to be efficient for large groups of 60 mice, as required in pre-clinical settings, utilising a short time window of 70 - 130 seconds per mouse per recording session.

**Figure 2.**
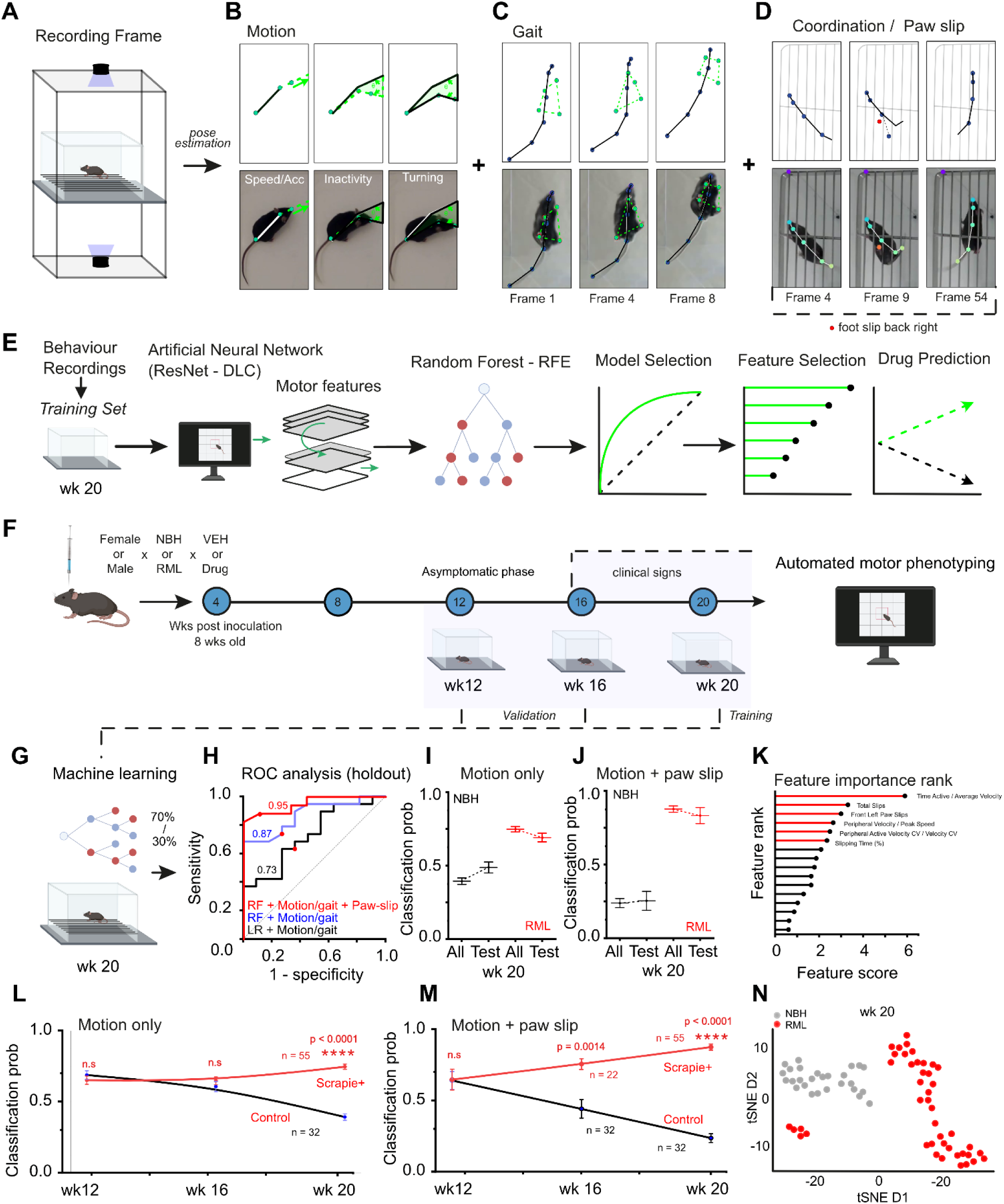
Pose estimation and machine learning can be used for automated classification of NBH and RML at 20 wks. (A) Recording setup for consistent imaging during each behaviour test for pose estimation using the Deeplabcut package. (B) Example of motion features. (C) Example of gait features. (D) Example of motor coordination and paw slip features. (E) Schematic of machine learning pipeline from pose-estimation coordinate data from 20 wk data using random forest recursive feature elimination (RFE), model comparison using ROC curves and drug comparisons on the best features. (F) Drug validation pipeline schematic with three time points for longitudinal validation where mice at 12 wpi are not expected to be symptomatic. (G) Data from training phase on 20-week data used for model training. (H) Comparison of linear models and random forest models on motion/gait and paw-slip data by ROC analysis. (I) Plot of classification probability for all mice vs the hold test set to assess overfitting for motion data. NBH and RML shown separately. (J) Plot of classification probability for all mice vs the hold test set to assess overfitting for motion + paw-slip data. NBH and RML shown separately. (K) Feature importance rank for the best model by RF-recursive feature elimination. (L) Plot of model (RF+Motion/gait) over time showing emergence of classification probability between control (n = 32) and scrapie inoculated (n = 55) in line with development of clinical signs. (M) Plot of model (RF+Motion/gait + paw slip) over time showing emergence of classification probability between control (n = 32) and scrapie inoculated (n = 55) in line with development of clinical signs. (N) TSNE plot illustrating differences between groups using the top performing features at week 20. Error bars are mean ± S.E.M. P values and n numbers are shown on the graphs unless < 0.0001 represented as ****. Statistics are corrected for multiple comparisons.

Using these recordings, we could track detailed movement of the head, body, and limbs using neural network-based estimation, implemented with the DeepLabCut package (Mathis *et al*., 2018). X/Y coordinate data was converted into motor features to describe the motion of each mouse in metrics such as speed, arena position, mobility time, and other rationally defined features (∼53). We then added combinatorial features for complex behaviour, including linear and non-linear combinations, to capture interactions (∼1325 without duplicates) that might reveal differences between control and disease-associated motor behaviour. After removing highly correlated features, we were left with a dataset of 94 such features, of which 60% were complex and the remaining “rational” or simple features.

Using this data, we then trained machine learning-based classifiers on control or scrapie inoculated mice and selected the models that best predicted disease status to enable disease-associated feature selection (Fig. 2E). Our aim was to generate a model that could be used to estimate the efficacy of a potential drug either directly or by prioritising features that best describe disease progression. To this end, we chose to record data across three time points of disease, from asymptomatic (12 wpi) to clinical signs (20 wpi), where only 20 wk data is used for training allowing for disease relevance of the model to be validated in previous timepoints in paired but asymptomatic mice (Fig. 2F). To ensure generalisation of the model, we trained it using a randomly selected subset (70%) of data from control or scrapie-injected mice at 20 wpi (Fig. 2G), the latest time point at which we recorded data, and then tested model performance on the remaining ‘holdout’ 30% of the data.

We tested a variety of models and assessed the ability of these models to correctly classify mice into control or scrapie groups using receiver operating characteristic (ROC) analysis (Fig. 2H). We reasoned that natural motor behaviour would be the most generalisable across treatment groups and disease stages, whereas paw-slip data would have specific utility for later stages of motor coordination loss in scrapie animals. Thus, we compared models using data from both combinations benchmarked against generalised linear model equivalents.

We found that a random forest model trained on motion-data alone had an ROC AUC of 0.87 and that including paw-slip data brought this value up to 0.95, indicating excellent discrimination that outperformed generalised linear models (AUC 0.73) trained on the same data (Fig. 2H). By comparing the classification probabilities for the scrapie and control groups with those from the holdout-test results, we were able to assess the extent of model overfitting. When trained on natural motion (Fig. 2I) or motion + paw-slip (Fig. 2J) data, with both models showing broadly consistent results, indicating good generalisation of the model to unseen data. To identify the most salient motor features extracted from recordings of open-field motion, and/or paw-slip, we employed recursive feature elimination in conjunction with the best random forest model to retain the motor features most informative about differences between control and scrapie mice at 20 wpi (Fig. 2K).

To further establish the selectivity of our model for features of scrapie that was established at data collected at 20 wpi, we next examined data collected from the same mice at 12 wpi during their asymptomatic phase, and at 16 wpi as an intermediate phase. We reasoned that this approach would reveal any non-disease specific differences between groups, prior to the presentation of motor signs (e.g. 18 wpi), that would affect classification. The best-performing models, trained on either motion and motion + paw-slip data, were applied to these timepoints and classification probability plotted longitudinally. These data strongly correlated with disease progression over time, starting with a mild skew towards the disease model over control (Fig. 2L). In the model including paw-slip data, we observed a stronger divergence, particularly in the control group, most likely because paw slipping is a rare event in these animals (Fig. 2M). The initial skew may have resulted from an imbalance in the data but did not alter classification. Lastly, the top performing features were used to visualise differences at 20 wpi using t-distributed stochastic neighbour embedding (tSNE) plots (Fig. 2N)

Overall, our model allows the selection of features that classify neurodegenerative progression that importantly generalises over time, successfully predicting disease absence during the asymptomatic phase. Furthermore, we found this method to be more sensitive than canonical methods at detecting early signs of disease.

### Pioglitazone has optimal chemical properties and neuroprotective potential for repurposing

We have previously shown that the safe and widely available anti-diabetic drug metformin has neuroprotective properties in RML scrapie mice (Harding et al., 2023). Here, we tested whether our preclinical analysis methods for scrapie progression could reveal neuroprotective effects of other candidate drugs. To select these candidates, we considered compounds approved for use in humans for metabolic disease, and that had chemical properties suggesting that they would have good solubility and oral bioavailability, and would likely cross the BBB to reach the CNS (Rankovic, 2015). Specifically, we extracted the chemical properties of candidate compounds from the ChEMBL database and calculated their multiparameter optimisation (MPO) scores. We considered MPO scores above 4 (out of 6) to indicate compounds with good brain accessibility (Wager *et al*., 2016). As our values relied partially on calculated properties from the ChEMBL database (Mendez *et al*., 2019), we validated the predictions against known published MPO values of ∼100 compounds. Where chemical features are known we predicted with complete accuracy (Wager *et al*., 2016). We then estimated compound solubility (LogS), using the general solubility equation (Ran and Yalkowsky, 2001), and considered scores between -2 and -4 to be sufficiently water soluble for oral availability. Specifically, we found that the anti-diabetic drugs pioglitazone and rosiglitazone have MPO scores of between 5.2 and 5.7 suggesting CNS penetrance and, have calculated LogS values around -3.5, are clinically available in tablet form, and have previously been associated with neuroprotective effects in humans and other model systems (Sato *et al*., 2011; Gupta and Gupta, 2012; Seok *et al*., 2019). Since rosiglitazone has been suspended from some markets, we selected pioglitazone for further study.

### Pioglitazone slows behavioural markers of scrapie in canonical and automated assessments of disease

To evaluate the anti-diabetic drug pioglitazone for neuroprotection over a long treatment time course, we added it to a mouse balanced control diet at 250 mg/kg for an estimated daily dose of 40 mg/kg/day for each mouse. We started treatment at 12 wpi, at the start of the behavioural studies and before the onset of clinical signs of prion disease, and continued treatment for the remaining time course of 8 weeks (Fig. 3A). Pioglitazone administration led to a modest increase in the body weight of both control and scrapie-inoculated groups (Supp. Fig. 2A-C), but all groups of mice reached similar mean body weights by 20 wpi.

**Figure 3.**
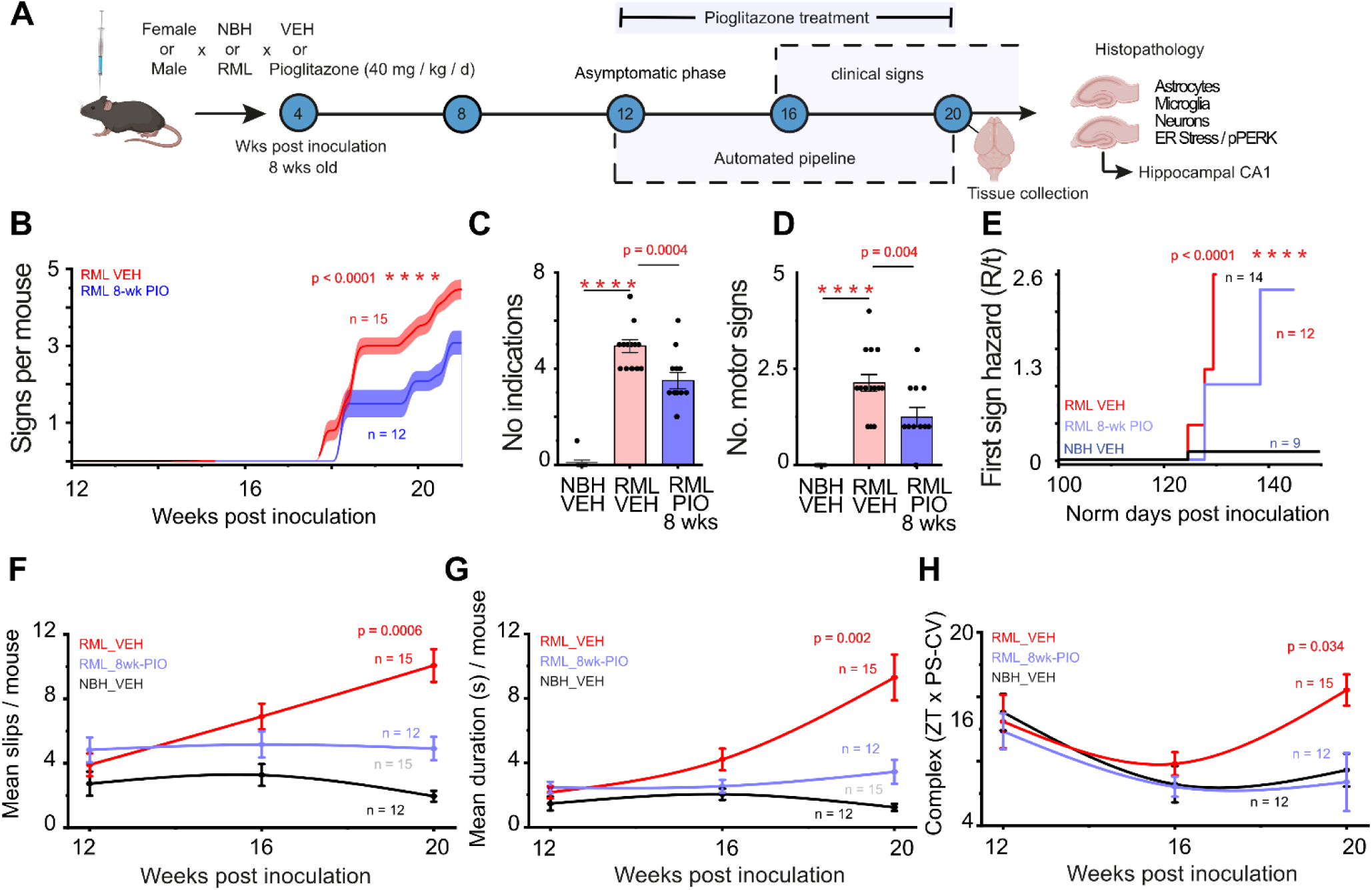
Pioglitazone shows evidence for neuroprotection in canonical and automated assessments of disease. (A) Schematic of drug testing pipeline with male and female mice. Highlighted areas show the time course for the automated pipeline vs classical clinical signs. Pioglitazone treatment at an average 40 mg / kg / day for 8-wks is highlighted above. After 20 wpi, mice used for behavioural analysis are culled for histological analysis. (B) Accumulation of all clinical signs and early indicators per mouse for each disease and drug treatment. (C) Total number of indications (signs + observations) of prion disease. (D) Total number of motor signs between disease and drug treated groups. (E) Time to first sign by hazard accumulation in normalised time. (F) Mean slips per mouse calculated from pose-estimation data over time. (G) Mean duration of slips over time between disease and drug treated groups. (H) Change in a complex movement feature over time between disease and drug treated groups. Error bars are mean ± S.E.M. P values and n numbers are shown on the graphs unless < 0.0001 represented as ****.

We next observed mice for clinical signs of scrapie (Fig. 3B) and found that treatment with pioglitazone substantially delayed their onset. We also found that the total number of indications (Fig. 3C), early signs combined with indicators of disease, were decreased, as well as the fraction of motor signs observed (Fig. 3D), Furthermore, the hazard rate to the development of the first observable sign was significantly longer in pioglitazone-treated groups (Fig. 3E). Overall, we found that pioglitazone treatment in RML scrapie-inoculated mice substantially slows clinical sign presentation.

To test if the results from subjective phenotyping for the appearance of clinical signs could be replicated and extended by automated behavioural phenotyping over the disease time course, we considered motor features that separated control and scrapie-injected animals (Fig. 2H). Specifically, we found that treatment with pioglitazone significantly reduced the number of paw slips by over 50% at 20 wpi and normalised the trajectory of motor coordination loss to approximately that of control animals (Fig. 3F). These findings were corroborated by a substantial decrease in the average duration of paw slips at both 16 wpi and 20 wpi, demonstrating that the beneficial effects of treatment are detectable substantially earlier than with classical observations (Fig. 3G). To corroborate these results, we found that a complex motor feature made from combinations of motor variables independent of paw-slip data, showed clear changes in disease trajectory over time (Fig. 3H). Complex features provide sensitive ways of capturing complex motor interactions without introducing conscious or unconscious bias about which features are likely to be experimentally important ahead of time. In all features shown, control (NBH) mice given pioglitazone were not different in weight from control (NBH) given vehicle treatment (Supp. Fig. 3A-C)

### Pioglitazone reduces markers of neuronal ER stress in scrapie mice

The action of pioglitazone in these animals suggested a neuroprotective mechanism in the brain. We have previously shown that the action of metformin, in rescuing motor function in neurodegeneration (Harding *et al*., 2023), appeared to work directly on deep brain structures. Similarly, other work has suggested a role of pioglitazone action directly on brain microglia that may aid in its neuroprotective properties (Ji *et al*., 2010; Zhao *et al*., 2016; Machado *et al*., 2019; Yeh *et al*., 2021).

To gain mechanistic insight into the neuroprotective effects of pioglitazone, we perfused scrapie-inoculated mice treated with this drug or vehicle at 20 wpi to enable histological analysis of the brain. Since pioglitazone has been previously shown to act on microglia via PPARγ (Ji *et al*., 2010; Machado *et al*., 2019), we immunostained brains for proteins enriched in microglia (Iba1) and astrocytes (GFAP) (Fig. 4A). To our surprise, there was little evidence of a difference in these markers (Fig. 4B & C). However, staining for the ER stress marker pPERK showed a strong reduction (Fig. 4D & E) consistent with our previous data for metformin. Gene expression data by qPCR showed no changes in *Cd68*, *Rbm3, Xbp1*, *Pikfyve* and a slight increase in *Atf6* (Supp. Fig. 4A-E). Together, these data suggest that pioglitazone may be achieving its neuroprotective effects by reducing neuronal stress rather than by reducing disease-associated neuroinflammation.

**Figure 4.**
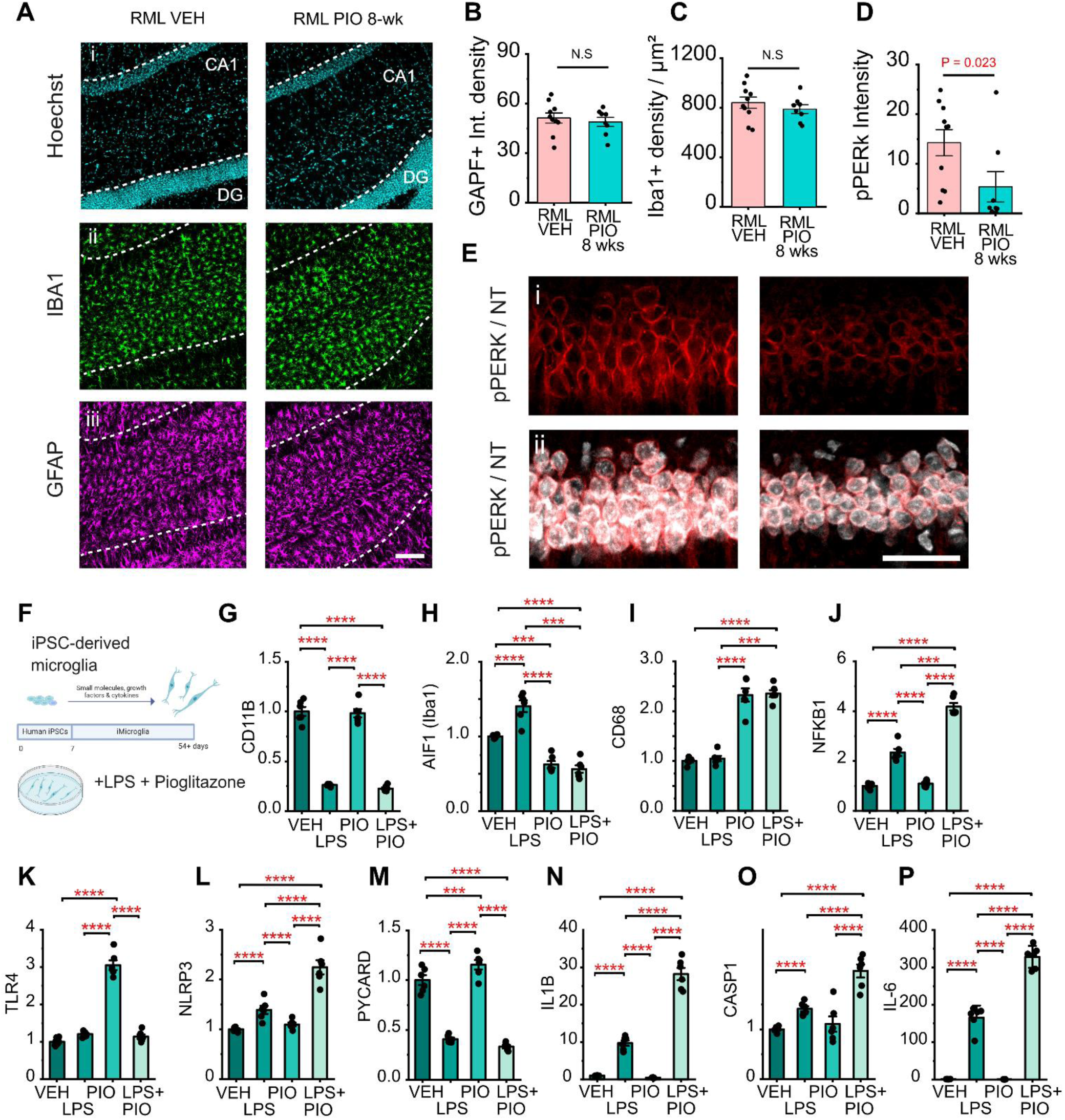
Pioglitazone reduces pPERK expression *in vivo* and microglial gene expression *in vitro*. (A) Representative hippocampal histology comparing the brains of RML scrapie inoculated mice treated with vehicle or pioglitazone, and stained with Hoechst, and immunohistochemistry for markers of microglia (Iba1) and astrocytes (GFAP). (B-D) No. significant differences were observed in the intensity of cells immunopositive for GFAP (B), or Iba1 (C) but a significant reduction in pPERK was seen between vehicle (n=10) or pioglitazone (n=8) treated mouse brains (E) Example of pPERK expression in hippocampal pyramidal neurons also labelled with NeuroTrace. (F) Schematic of microglial differentiation from human iPSCs before treatment with vehicle or LPS (100 ng / mL) and vehicle or pioglitazone (20 μM). (G-P) Expression levels of the indicated genes, as quantified by RT-qPCR. n=6 replicates/condition. Error bars are mean ± S.E.M. P values and n numbers are shown on the graphs unless < 0.0001 represented as ****.

### Pioglitazone cell-autonomously alters microglial gene expression

Since pioglitazone has previously been described to act on microglia (Ji *et al*., 2010; Machado *et al*., 2019), but we saw little evidence of this in our histological analysis in prion mice, we set out to directly test its potential cell-autonomous effects on glia to resolve this apparent discrepancy. To the end, we differentiated human iPSCs into microglia using established protocols (Washer *et al*., 2022), and exposed them to either vehicle or pioglitazone (Fig. 4F). Furthermore, we tested whether pioglitazone could affect microglial cell state when stimulated with the potent proinflammatory stimulus lipopolysaccharide (LPS). We found that, pioglitazone treatment did not alter expression of *CD11B* (*ITGAM*) alone, or affect its response to LPS (Fig. 4G). However, pioglitazone was able to significantly inhibit the LPS-induced upregulation of *AIF1 (IBA1)* (Fig. 4H), which is known to increase following microglial activation and plays an important role in phagocytosis (Streit *et al*., 2009). We observed an upregulation of *CD68*, a commonly used marker of functionally activated microglia, following pioglitazone treatment (Fig. 4I), consistent with reports that PPAR-γ activation promotes microglial phagocytosis (Krishna *et al*., 2021). We also noted large changes in cytokine and inflammasome-related genes in response to LPS treatment as expected, and found that pioglitazone treatment was unable to inhibit these changes and, in some cases, accentuated them (Fig. 4J-P), though the strong LPS stimulus may have obscured more physiologically relevant responses. Overall, we conclude that pioglitazone is able to act directly on human microglia to modify gene expression in both the un-stimulated and LPS-stimulated state, suggesting microglia may be involved, at least in part, in the neuroprotective effects we observed.

## DISCUSSION

We have shown that preclinical testing of candidate drugs in mice can be automated using pose-estimation and machine learning to provide an unbiased assessment of drug efficacy and potentially greater sensitivity. We then show neuroprotective potential for pioglitazone in a scrapie model of aggressive neurodegeneration which we show reduces markers of neuronal ER stress. Here, we discuss our findings and the limitations and potential of our study in greater detail.

Selecting the appropriate animal model for preclinical testing is essential for interpreting phenotypes and increasing the likelihood that findings in animals will translate to humans. To model neurodegenerative disease, we therefore selected to work with RML scrapie since this model: 1) phenocopies human prion disease without genetic manipulation 2) includes the full spectrum of pathological features common to many human neurodegenerative diseases including progressive synaptic loss, ER stress, neuroinflammation, and neuronal loss that are incompletely captured in other transgenic models (Richardson and Burns, 2002; Mallucci, 2009; Freeman and Mallucci, 2016), 3) is versatile since RML brain homogenate can be introduced on different genetic backgrounds, in different brain regions, and at different time-points, and 4) is relatively aggressive and shows stereotyped progression, allowing for neuroprotective effects to be determined with greater confidence over a shorter period of time than in alternative animal models (Yoshiyama *et al*., 2007). RML scrapie is sometimes injected in transgenic animals overexpressing prion protein to further accelerate the disease time course (Watts and Prusiner, 2014), but here we selected to work with young adult male and female C57BL/6J mice to leverage the wealth of phenotypic data available on this genetic background and age group.

To characterise RML scrapie progression in this model histologically, we focused our molecular characterisation on the hippocampal CA1 region of these mice, since this brain region is involved in memory impairments seen in common neurodegenerative diseases such as Alzheimer’s Disease. This brain region is also physically near the brain homogenate inoculation sites in the parietal cortex, allowing the early histological changes associated with scrapie disease progression to be detected. We found a stereotyped time course of histopathological features, starting with an increase in markers of reactive astrocytes with small increase in the ER stress as measured by pPERK at 12 wpi, during the asymptomatic phase of disease. This was followed by a sustained microglia activation and a further six-fold increase in ER stress by the clinical phase of 20 wpi, coinciding with neuronal loss. These significant and progressive histopathological hallmarks can be exploited for mechanistic understanding of disease, using longitudinal study designs that examine the effects of drug treatments on the molecular progression of disease.

Survival assays have been a bedrock method in prion research for many decades and have been successfully used to reveal differences in infectivity between prion strains, the importance of species barriers in disease transmission, and the effectiveness of decontaminating agents (Taylor, 1999; Sakudo, Anraku and Itarashiki, 2020). However, confirmatory motor signs appear late in prion disease time course, so there is a clear advantage to have earlier indicators of disease progression that shorten experimental paradigms. In addition, we and others have observed bladder complications in some strains of female mice injected with scrapie that arise before confirmatory motor signs that complicates the use of both sexes in experiments where later disease stages must be reached for the assessment of clinical signs.

To complement and extend traditional histological analysis and subjective scoring of behavioural signs of prion disease, we developed methods to objectively and longitudinally measure motor behaviours over the course of RML scrapie disease progression. We then employed a pose estimation and machine learning-based behavioural analysis pipeline to attempt to replace many scrapie clinical observations. Our aim was to produce a pipeline that would be inherently unbiased but at the same time would reduce the extensive labour of standard behaviour tests, reduce batch effects from being able to use larger well powered group sizes, allow frequent mixed sex groups, and have greater sensitivity due to the use of new computational tools.

This innovation has several substantial benefits that increase the utility of the model. First, data collected and analysed in this manner are objective and therefore provide greater consistency than can be easily achieved between users or laboratories. For example, adding drugs to food or water can change its texture and/or appearance and be unintentionally unblinded. This method removes much of the challenge of blinding during experimentation as the scientist has little influence over the outcome and focuses instead on recording consistency.

Second, the use of standardised arenas, and standardised experimental design and recording methods, facilitates replication across experimental batches and across research groups. Third, the resulting behavioural data are rich in features and can readily be re-analysed by other groups or by ourselves as analysis methods improve. Fourth, we found clear behavioural differences in prion disease several months before the appearance of confirmatory motor signs, enabling the inclusion of female mice that would otherwise have been excluded due to complicating bladder issues. We found that these early motor signs were predictive of scrapie disease progression in our previous work (Harding *et al*., 2023) and validated again here by the progressive accumulation of motor signs over time in an automated system. Finally, collecting longitudinal data allows neurodegenerative disease progression to be followed in individual mice, providing greater confidence that treatment effects on disease progression are consistent despite inevitable inter-individual variability.

We selected to test pioglitazone in this model system, since this drug is widely used around the world for the treatment of diabetes with clearly understood safety and therapeutic profile, and since it has previously been shown to have neuroprotective properties in other mouse models of neurodegenerative disease (Ji *et al*., 2010; Seok *et al*., 2019). Furthermore, we previously showed that metformin has direct neuroprotective properties in RML scrapie mice even at concentrations as low as ∼250 nM (Harding *et al*., 2023), and both pioglitazone and metformin are used for the treatment of diabetes (frequently in combination), albeit via distinct mechanisms with metformin acting (at least in part) via AMPK activation and pioglitazone via PPARγ activation. We found that RML scrapie mice treated with either metformin (Harding *et al*., 2023), or pioglitazone had a reduced rate of disease progression as assessed by slower acquisition of clinical signs of prion disease, and also reduced PERK phosphorylation in CA1 hippocampal pyramidal neurons. These findings suggest that both metformin and pioglitazone reduce neuronal ER stress and slow neurodegenerative disease progression, though the precise mechanisms are unclear. In this study, we extend these findings by showing that pioglitazone significantly slowed disease progression as determined by our unbiased preclinical behavioural pipeline, providing greater insight into the time frame and behavioural features of these neuroprotective effects, and providing a framework for future drug testing.

To gain insight into the associated molecular and cellular features accompanying reduced neuronal ER stress and slower disease progression in response to pioglitazone treatment, we carried out histological analysis for markers for neuroinflammation. Previous studies have suggested that pioglitazone may act to suppress microglial activation (Ji *et al*., 2010; Zhao *et al*., 2016), and metformin might act via cell-autonomous effects on microglia and astrocytes to suppress scrapie-associated neuroinflammation. We see consistency between the ability of metformin and pioglitazone to both reduce pPERK immunoreactivity and to delay motor symptoms. We did not observe significant differences in the immunoreactivity of microglial or astrocytic markers in the brains of vehicle or pioglitazone-treated mice. This was despite pioglitazone’s clear effect on expression of *Cd11b*, *Aif1* (*Iba1*) and *Cd68*, when iPSC-derived microglia were challenged with LPS (Fig. 4). This suggests that reduced neuronal ER stress may be a common neuroprotective mechanism. However, the divergent effects on microglia and astrocytes between these treatment groups, suggests that neuroinflammation, at least how we define it histologically, is not as important an indicator of the state of disease progression or drug mechanism of action.

While further studies are needed to explore likely mechanisms of action, these findings suggest that pioglitazone is unlikely to reduce neuronal ER stress primarily by reducing neuroinflammation, suggesting either cell-autonomous protective mechanisms on neurons, or systemic effects such as improved glucose homeostasis that then support neuronal health and survival. Understanding these aspects are key to moving drugs like these into successful trials as the mechanism is a key part of appropriate clinical trial design. We note that largest study of pioglitazone for human dementia treatment we are aware of “TOMORROW” (Burns *et al*., 2021), used very low dose pioglitazone (0.8 mg /day vs 40 mg / day) in healthy patients stratified by predicted risk. This study also faced other hurdles, without the use of biomarkers to enrich the patient cohort, and a change in futility end points during the study. We believe that better mechanistic understanding of these therapies will aid improve clinical trial design as well as the stratification of patients into cohorts most likely to benefit from treatment.

### Limitations

There are some clear limitations to our study. We built it around a single neurodegenerative model, but the use of automated pre-clinical phenotyping has been designed to generalise to any model with mild or greater motor symptoms (e.g. PS19 mice, Yoshiyama et al., 2007) and could help automate and systematise pre-clinical drug testing. The use of machine learning and ensemble models in animal research is always a challenge due to the requirement for large group sizes. To address this challenge, we generated control groups of 82 mice in our first iteration, but recognise that model performance will increase as animal number increases in future studies. Indeed, such a dataset could also provide a more generalisable model that other groups can benchmark against and ultimately produce shareable datasets that make animal experiments more efficient and robust. We analysed behaviour in thousands of tests performed at three discrete time points. We recognise that additional time-points and types of behavioural analysis (e.g. cognitive tests) would improve our understanding of disease progression. Looking forward, 3D pose-estimation methods (Lauer *et al.,* 2022), as well as 24-hour recording in home cages would further enhance our ability to collect and rigorously analyse pre-clinical data.

The scrapie model is not commonly used in neurodegenerative disease research, although requires only an inoculation of misfolded PrPsc into wild-type animals, and since it recapitulates bona fide neuronal loss preceded by progressive histopathological features common to many human neurodegenerative diseases that also have protein aggregation and ER stress as likely shared disease mechanisms. Although there are prion disease has distinct features as a proteinopathy (Meisl *et al*., 2021), misfolded protein are common to Alzheimer’s and Parkinson’s and both amyloid and tau pre-formed fibrils can seed and propagate in the brains of mice and humans (He *et al*., 2018; McAllister *et al*., 2020; Condello *et al*., 2023), even in cases of Alzheimer’s (Banerjee *et al*., 2024). We view misfolded protein as the first event in a cascade that causes neurodegeneration across multiple types of dementia, and identifying compounds that slow prion disease may therefore reveal neuroprotective agents that cut across neurodegenerative disease types.

We propose that the mouse scrapie model and the methods we developed here provide unbiased and sensitive assessment of pre-clinical drug efficacy that can accelerate the rate of drug testing, and increase the likelihood that discoveries made in this model will translate into successful human clinical trials.

## MATERIALS AND METHODS

### Animals and colony maintenance

Animal work was performed under a Project Licence (PPL7597478) administered by the UK Home Office. University of Cambridge Animal Welfare and Ethics Review Board (AWERB) approved all procedures and protocols adhered to 3Rs and ARRIVE guidelines. All mice used in this study were of the C57BL/6J strain and were purchased from Charles River Laboratories (Saffron Walden, UK). Upon arrival, mice were fed a standard chow diet and group housed at 3 or 5 mice per cage in individually ventilated cages for seven days to acclimatise them to the specific pathogen free facility. The animal facility was maintained on a standard 12-hour light/dark cycle and temperature was controlled at 22 ± 2°C. Water and food were available *ad libitum*.

### Inoculation with brain homogenates

C57BL/6J male and female mice were acclimatised to the facility for one week in cages of three, and then randomly allocated for inoculation with either Rocky Mountain Laboratory (RML) prion or Normal Brain Homogenate (NBH) between the ages of 4 and 6 weeks. Mice were inoculated by injection in the parietal cortex with 30 µl of a 1% homogenate solution (gift from the laboratory of Prof. Giovanna Mallucci), while under isoflurane anaesthesia as previously reported (Halliday *et al*., 2017). Mice were monitored until recovered and then returned to their home cage.

### Diet and treatment group randomization and administration

All mice were fed standard chow diet until 7 weeks old, before acclimatisation to balanced control diet (10% fat diet D12450Ji, Research Diets). Mice were weighed at least once weekly. Scrapie-inoculated and control mice were maintained on their allocated diets for approximately 21 weeks post inoculation before culling at a fixed time point. In the week prior to culling, mice were provided with wet diet supplementation including M31 diet gel and hydrogel during the weight loss phase of the disease. Control mice were provided the same supplementation. Pioglitazone was added to control diet food (D23042701i) as purchased at 250 mg/kg diet for an average dose per mouse of 40 mg/kg/day.

### Terminal tissue and blood collection

At the time of culling, a heparin-coated (375095, Sigma Aldrich) syringe with a 23G blunt-end needle was used to collect blood by cardiac puncture while under deep terminal anaesthesia, followed by cervical dislocation. Blood was transferred to tubes containing lithium heparin (450537, Greiner Bio-One MiniCollect™) and plasma was separated by centrifugation at 800 x g. Organs were then immediately dissected, snap-frozen on dry ice, and stored at -70°C. Brains were divided in two along the midline, with half micro-dissected into different brain regions and snap frozen as described above, and half reserved for histology by overnight fixation in 4% PFA followed by washes in PBS and storage in 30% sucrose the following day.

### Immunohistochemistry

Prion-inoculated brains that had been fixed in 4% PFA overnight were decontaminated by complete immersion in >95% formic acid for at least 60 mins at room temperature, before washing in PBS and re-fixation in 4% PFA. For immunohistochemistry, 25-µm thick coronal brain sections were obtained using a Leica VT1000S vibratome (Leica Biosystems, Germany). Then, free-floating sections were subjected to antigen retrieval (Zur *et al*., 2024) in citrate buffer (pH 6) at +90°C (for pPERK staining) or autofluorescence quenching in 0.3% Sudan Black B (Oliveira VC, 2010) in 70% ethanol for 20 min (for Iba1/GFAP staining). Brains were treated with a blocking solution containing 10% NDS (Jackson ImmunoResearch Labs, USA), 1% BSA (Miltenyi Biotec Ltd.), and 0.3% Triton X-100 (Sigma-Aldrich, USA) for 1 hour at room temperature, and then incubated overnight at +4°C with primary antibodies: rabbit anti-Iba1 (1:750, Abcam, UK), chicken anti-GFAP (1:1500, Antibodies.com, UK), rabbit anti-NeuN (1:1000, Merck Millipore, USA), or rabbit anti-pPERK (1:250, Cell Signaling Technologies, USA). After washing with PBS, the sections were incubated with donkey anti-rabbit AF488, goat anti-chicken AF647, and donkey anti-rabbit AF555 secondary antibodies (1:1000; Invitrogen, USA) for 1.5 hours at room temperature in the dark. Samples were then rinsed in PBS, counterstained with NeuroTrace 640/660 (1:100; Thermo Fisher Scientific, USA) and Hoechst 33342 (1:10000; Invitrogen, USA), placed on histological slides, and mounted with Aqua/Poly-Mount medium (Polysciences, USA). Images of the hippocampal CA1 area were taken using a Leica SP8 confocal microscope (Leica Biosystems, Germany) at 20× or 60× magnification.

### Clinical observations

Mice were assessed for prion disease by the development of early signs as previously described (Moreno *et al*., 2013; Halliday *et al*., 2017; Smith *et al*., 2020), and for additional indicators of loss of motor coordination (Harding *et al*., 2023). Prion observations were blind to any previous records of early signs, but could not be blinded to treatment variables as drug presence in the food was visually apparent. Normally, the presence of one early sign and a confirmatory clinical sign are sufficient to diagnose late-stage prion disease, leading to culling the mouse. Early signs such as motor coordination in canonical methods were classified on the observation of paw slips. Specifically, Motor 1 (M1) was classified as any obvious foot slip on bars of the standard IVC homepage.

Inoculated animals are known to be susceptible to bladder enlargement with female mice being particularly prone to this complication. All mice showing indicators of illness were first assessed for bladder enlargement and all mice were checked on autopsy for presence of gross pathology, including blood around the organs, bladder enlargement, or features of liver disease. On rare occasions mice with ‘large’ enlarged-bladders, detectable through the abdomen, presented with an inability to urinate and hunched posture. These mice were culled, confirmed to have bladder complications on autopsy and excluded from survival data (<5% mice).

### Pose estimation

Pose estimation was performed using the DeepLabCut Package for single animals. Artificial neural networks using a pre-trained Resnet-50 were updated and trained on the University of Cambridge high performance cluster using training set images from our experimental arena. Separate networks were trained for motor, gait and paw-slip assays. Frames for labelling were selected by k means clustering with 4-6 frames per video and training for 400,000 iterations. For motor analysis, 152 frames were used with 8 for testing, resulting in a p-cutoff error of 1.88 pixels. For gait analysis 254 frames were used, with 15 for testing resulting in a p-cutoff error of 6.01 pixels. For paw-slip detection frames selection was biased to RML data to increase slip numbers in the training set and slip variety. Frames were also selected using k-means clustering with additional frames showing various paw slips included through manual selection, resulting in 256 used for training with 14 frames used for testing resulting in a p-cutoff of 5.49 pixels.

### Machine Learning

XY coordinates from pose estimation of tracked body points were generated for three sets of behavioural features, locomotion (motor) features, paw slip features, and gait features. Missing (NA) values were interpolated using mice (3.16.0) in R with predictive mean matching, performing 5 imputations and 5 iterations, all with a constant seed value. Feature engineering incorporated domain-specific knowledge to create new features, as well as the generation of interaction and ratios of existing features. The three sets of behavioural features were first checked for non-zero variances. Data was partitioned into training and testing sets (70 / 30), which were standardised separately. Pearson correlation analysis was performed to remove highly correlated features (r > 0.85). Two classification models-logistic regression and random forest - were developed and evaluated using 5-fold repeated cross-validation (n=3). To address multicollinearity, features were iteratively removed in logistic regression based on the variance inflation factor (<5), and random forest employed recursive feature elimination. Following hyperparameter tuning, the following was used for the respective models: a) - motor: mtry was 1 and ntree was 100, b) motor + gait: mtry was 6 and ntree was 100 and c) paw slip + motor: mtry was 5 and ntree was 50. Modelling was performed in R using packages tidyverse (2.2.0), caret (6.0-94), and ggplot2 (3.5.1).

### Human iPSC-microglia

Human iPSC-microglia were generated based on the previously published protocol (Washer *et al*., 2022). Briefly, iPSCs were cultured in Geltrex coated 6-well plates in OxE8 medium for maintenance. Following 1-2 passages, iPSCs were detached and collected as a single cell suspension. After centrifugation, 4 million cells were plated into Aggrewell 800 plates in 2 mL EB Induction medium (OxE8 medium + stem cell factor (SCF) + bone morphogenetic protein 4 (BMP4) + vascular endothelial growth factor (VEGF)) per well to generate embryoid bodies (EBs). To allow the formation of EBs, cells remaining in AggreWell plates received daily 75% media change for 6 days. After 6 days, EBs were then harvested and equally distributed to two T175 flask containing 18 mL of EB Differentiation medium (X-Vivo 15 medium + Glutamax + β-mercaptoethanol + interleukin 3 + macrophage colony-stimulating factor (M-CSF). The EBs were kept in EB Differentiation medium at 37°C and 5% CO2 with full media changes every 7 days. After 2-3 weeks, non-adherent microglial precursor cells (PreMac) were released into the medium from EBs. PreMacs were harvested and strained through a 40 µm cell strainer. Harvested PreMacs were pooled and sustained in EB Differentiation medium in T75 flasks with weekly media changes. Once sufficient cell numbers were collected, PreMacs were seeded onto 6-well plates at a density of 100,000 cells/cm^2^ in maturation medium (advanced DMEM-F12, Glutamax, penicillin-streptomycin, IL-34, granulocyte-macrophage colony-stimulating factor (GM-CSF), transforming growth factor beta 1 (TGFb1), and M-CSF) to differentiate and mature PreMacs to iMicroglia for 14 days. The iMicroglia were further matured for 14 days in maturation medium prior to experimentation. Matured iPSC-microglia was treated with either Pioglitazone (20 µM) or Vehicle (DMSO) 1 hour prior to vehicle or LPS (100 ng/ml) challenge for 24 hours.

### RT-qPCR

Total RNA from treated iPSC-microglia was extracted using the RNeasy Plus Mini Kit (Qiagen) and half of the hippocampus from inoculated mice treated with vehicle or pioglitazone was extracted using the RNeasy Plus Universal kit (Qiagen) according to the manufacturer’s instructions. Extracted total RNA (0.6-1 µg) was reverse transcribed into cDNA using the iScript^TM^ reverse transcription supermix. Target genes of interest were determined using the Taqman gene expression ‘assay-on-demand^TM^’ assays (Applied Biosystems, Foster City, CA) using FAM-labelled probes for target genes and *ACTB* (Hs01060665_g1) and *GAPDH* (Hs99999905_m1) as reference genes. The primer/probe targets were *ITGAM* (Hs00167304_m1), *AIF1* (Hs01032552_m1), *CD68* (Hs02836816_g1), *NFKB1* (Hs00765730_m1), *TLR4* (Hs00152939_m1), *NLRP3* (Hs00918082_m1), *PYCARD* (Hs01547324_gH), *IL1B* (Hs01555410_m1), *CASP1* (Hs00354836_m1), and *IL6* (Hs00174131_m1). For the mouse experiments, *Actb* (Mm02619580_g1) and *Gapdh* (Mm99999915_g1) were used as reference genes, and primer/probe targets were *Cd68* (Mm03047343_m1), *Atf6* (Mm01295319_m1), *Xbp1* (Mm00457357_m1), *Rbm3* (Mm00812518_m1), and *Pikfyve* (Mm01257047_m1). The reactions were dispensed with Echo 525 Acoustic Liquid Handler (Beckman Coulter) and run on the QuantStudio™ 5 Real-Time PCR System in 384-well plates (Thermo Fisher Scientific). All expression assays were normalised to the geometric mean of the reference genes *GAPDH/Gapdh* and *ACTB/Actb f*or human/mouse experiments, respectively.

### Data Analysis

Data was analysed in either Origin Pro or R using custom scripts. Time to event data including survival was analysed using the Kaplan Meyer plot and log rank test. Two group analysis was performed by two-way t-test (equal or unequal variance) or Mann Whitney U, three levels by one-way ANOVA and two-factor tests analysed by two-way ANOVA. Variance was determined by the F test. The Bonferoni-holm method was used for correction of multiple comparisons. Image analysis was performed in Fiji ImageJ 1.54f using custom macros for measuring Iba1^+^ (MaxEntropy Threshold) and NeuN^+^ (Huang Threshold) cell densities, as well as Iba1^+^ microglial soma area and circularity. The “Integrated Density” function was used to analyse GFAP^+^ and pPERK^+^ images after applying a fixed threshold.

## Author Contributions

**E.C.H., H.-J.C.C** and **F.T.M.** designed experiments and drafted the manuscript with input from all authors. **E.C.H** analysed data and produced visualisations and performed experiments together with **H.-J.C.C**, **D.S**^1,3^**, C.R, S.O.Z, D.S**^4^**, I.M, J.J.C, S.J.W,** and **N.S. A.R.B** supported experiments and provided reagents. **F.T.M** acquired funding and assisted in data interpretation. **F.T.M** and **E.C.H.** conceived of and managed the project and personnel.

## Acknowledgments

We are grateful to Giovanna R. Mallucci for providing RML inoculum for experiments. F.T.M. is a New York Stem Cell Foundation - Robertson Investigator (NYSCF-R-156) and is supported by the Wellcome Trust and Royal Society (211221/Z/18/Z) and a Ben Barres Early Career Acceleration Award from the Chan Zuckerberg Initiative’s Neurodegeneration Challenge Network (CZI NDCN 191942) which supported this work and financially supported E.C.H, H.-J. C.C., C. R., D.S^1,3^. Seed grants to E.C.H were provided by the Cambridge Centre for Data-Driven Discovery and the Accelerate Programme for Scientific Discovery, made possible by Schmidt Sciences, LLC as well as The Alan Turing Institute. We thank the Disease Model Core (DMC) for assistance with mouse phenotyping. We thank the IMS Genomics and Bioinformatics Core (GBC) Facility for equipment support. DMC and GBC are funded by the UK Medical Research Council (MRC) Metabolic Disease Unit (MRC_MC_UU_00014/5), a Wellcome Trust Major Award (208363/Z/17/Z), and a Wellcome DRP. For the purpose of open access, the authors have applied a CC-BY public copyright licence to any Author Accepted Manuscript version arising from this submission.

---

**Supplementary Figure 1.**
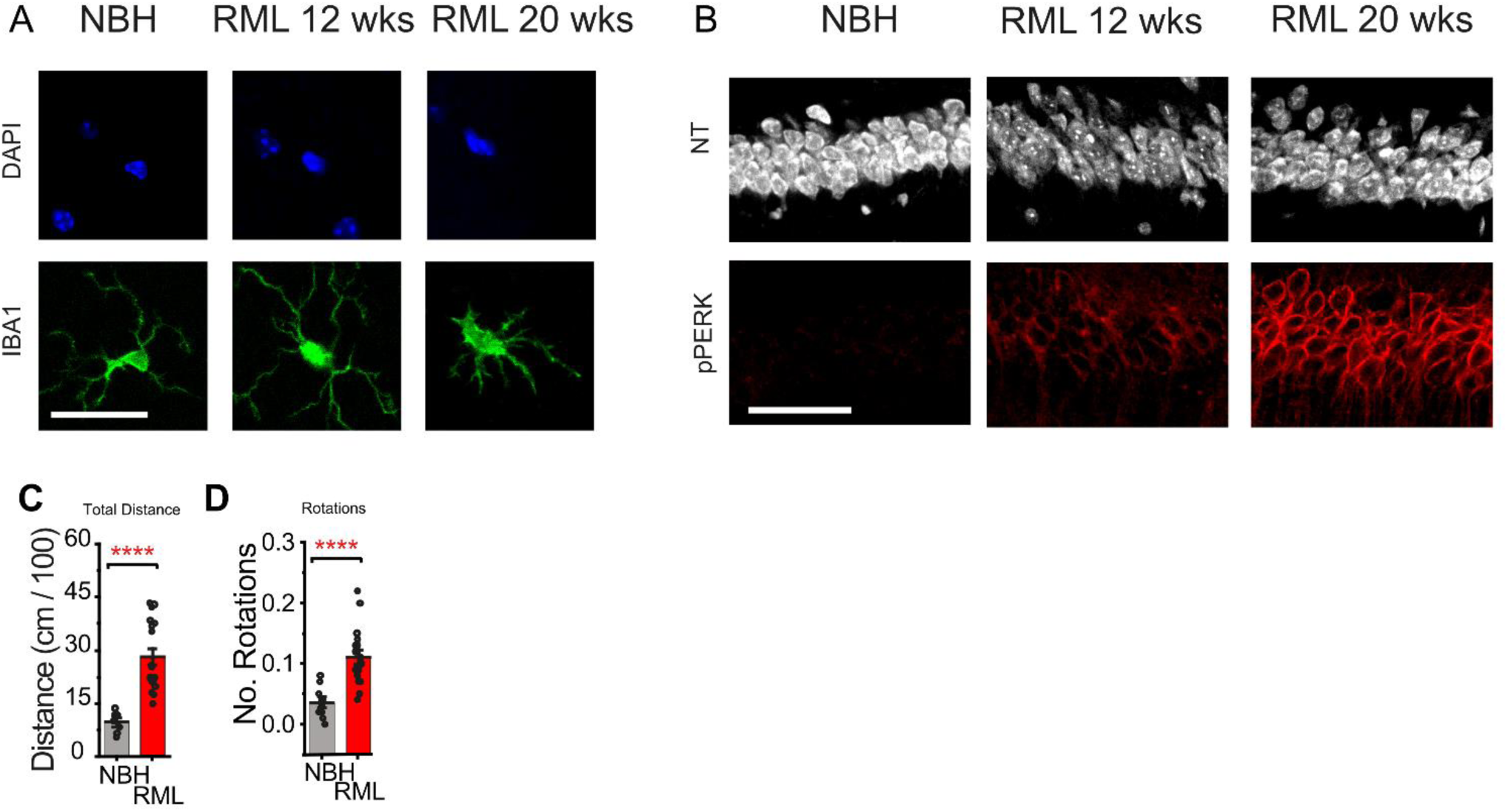
Microscopy images from brain histology and motor data. (A) High magnification images of microglia from the brains of control of RML-inoculated mice counterstained with DAPI and visualised by Iba1 immunoreactivity. Scale bar represents 20 μm (B) NeuroTrace staining and pPERK immunoreactivity. Scale bar represents 50 μm (C) Total distance moved in a locomotion experiment between control (n = 9) and scrapie (n = 18) mice at 20 wpi. (D) Total number of rotations between control (n = 9) and scrapie (n = 18) mice at 20 wpi. Error bars are mean ± S.E.M. P values and n numbers are shown on the graphs.

**Supplementary Figure 2.**
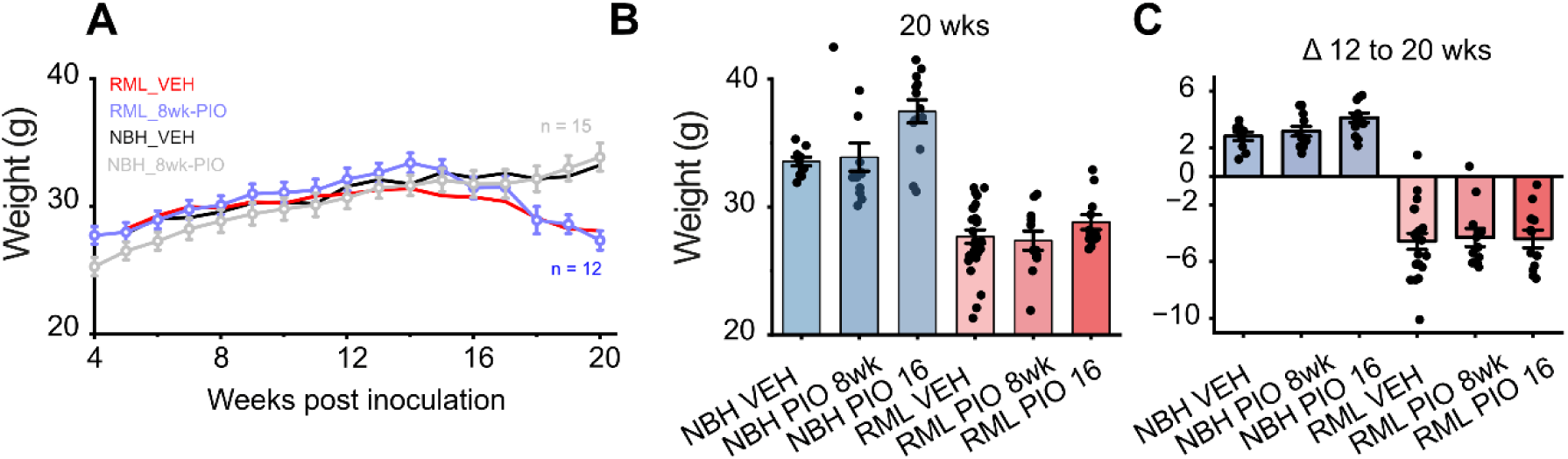
Body weight of mice during the experimental time course. (A) Body weight per mouse shown over time from 4 to 20 wpi. RML_VEH and NBH_VEH also shown in Figure 1B duplicated here for visualisation without error bars. (B) Absolute body weight at 20 wpi for each group. (C) Body weight changes from 12 wpi to 20 wpi. Error bars are mean ± S.E.M. P values and n numbers are shown on the graphs.

**Supplementary Figure 3.**
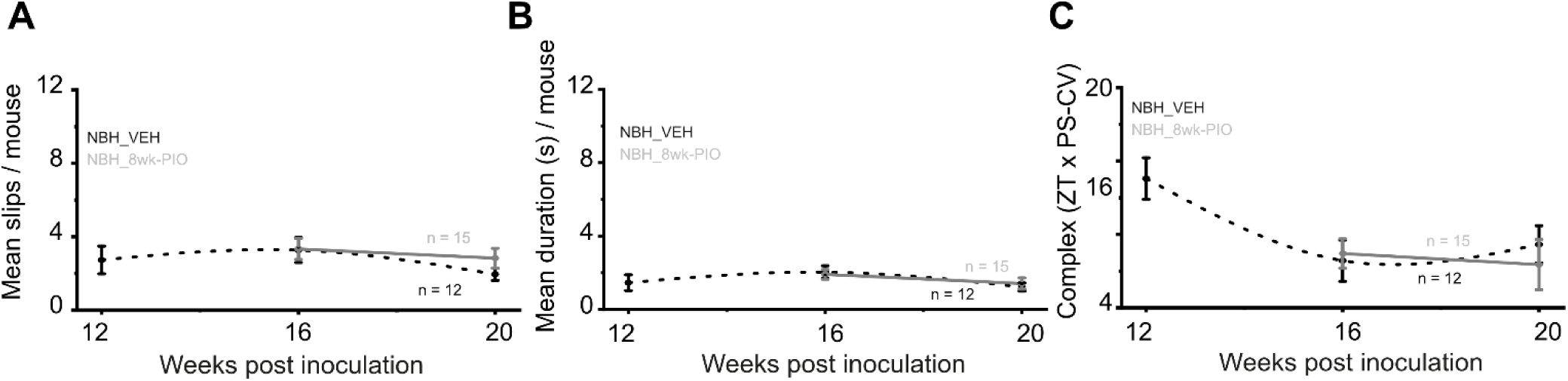
Pose-estimation data for control vs control with pioglitazone. (A) Mean number of paw-slips per mouse (B) duration of paw-slips per mouse calculated from pose estimation data over time for control vs control with drug treatment. (C) Change in complex moment feature over time from pose estimation data over time for control vs control with drug treatment. Error bars are mean ± S.E.M. P values and n numbers are shown on the graphs.

**Supplementary Figure 4.**
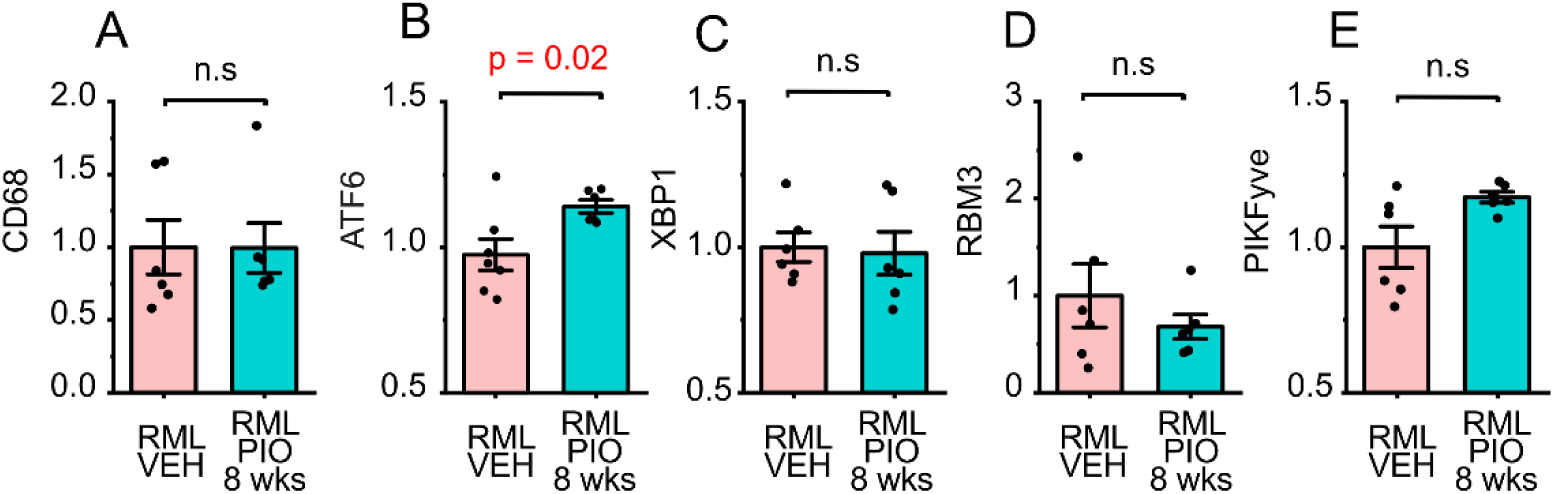
Pioglitazone-induced changes in gene expression in scrapie mice. Hippocampal samples collected at 20 wpi from RML-inoculated mice treated with vehicle or pioglitazone used to determine relative gene expression for A) *Cd68*, B) *Atf6*, C) *Xbp1*, D) *Rbm3* and E) *Pikfyve*. Error bars are mean ± S.E.M. P values are shown graphs for significant results.

## Notes

### Competing Interest Statement

The authors have declared no competing interest.

